# Expansion microscopy reveals insulin granule clustering in human β-cells in type 2 diabetes

**DOI:** 10.64898/2026.05.05.722840

**Authors:** Licia Anna Pugliese, Valentina De Lorenzi, Gianmarco Ferri, Ha Vo, Allison Lindquist, Marta Tesi, Carmela De Luca, Mara Suleiman, Lorella Marselli, Yongxin Zhao, Piero Marchetti, Fabio Beltram, Francesco Cardarelli

**Author notes:** equal contribution (Licia Anna Pugliese, Valentina De Lorenzi).

## Abstract

**Aims/hypothesis:** Quantitative nanoscale analysis of insulin secretory granules (ISGs) in human pancreatic tissue has been limited by the lack of imaging methods that combine high resolution with large-scale sampling. We aimed to establish expansion microscopy (ExM) as a platform for in situ, quantitative analysis of ISG organisation in human β-cells and to assess whether type 2 diabetes (T2D) is associated with alterations in granule size, abundance or spatial organisation.

**Methods:** We applied Magnify ExM to PFA-fixed, paraffin-embedded pancreatic tissue sections from 6 human donors, 3 non-diabetic (ND) and 3 T2D, enabling super-resolution optical imaging of insulin-labelled granules. Insulin-positive structures were segmented and analysed using a morphometric pipeline to quantitatively assess size, shape and spatial features. Granule clustering was quantified based on combined area and roundness criteria.

**Results:** The diameter distribution of highly circular granules was similar between ND and T2D samples and estimates of granule number per cell indicated only a modest reduction in T2D (∼25%). In contrast, mapping insulin-positive structures in a roundness–area space revealed a marked enrichment of large, irregular objects consistent with granule clustering in T2D. The fraction of clustered granules was significantly increased in T2D and strongly inversely correlated with insulin stimulation index (r = −0.85).

**Conclusions/interpretation:** These results establish expansion microscopy as a powerful platform for quantitative nanoscale analysis of human pancreatic tissue and identify altered spatial organisation of insulin granules, rather than marked granule depletion, as a prominent feature associated with β-cell dysfunction in T2D.

**Research in context:** *What is already known about this subject?:*

- β-cell dysfunction in type 2 diabetes is often attributed to reduced insulin content or β-cell loss.
- Insulin secretory granules (ISGs) have been characterised ultrastructurally, but quantitative analysis in human tissue remains limited.
- Super-resolution approaches, including expansion microscopy, are emerging tools for nanoscale imaging in biological tissues.

*What is the key question?:*

- Is β-cell dysfunction in type 2 diabetes associated with depletion of insulin granules or with altered spatial organisation?

*What are the new findings?:*

- Insulin granule size distribution is largely preserved in type 2 diabetes, with only a modest reduction in granule number per cell.
- A significant increase in insulin granule clustering is observed in diabetic β-cells.
- Granule clustering is strongly inversely correlated with insulin secretion in the same donor tissues.

*How might this impact on clinical practice in the foreseeable future?:*

- Identifying altered granule organisation as a feature of β-cell dysfunction may help refine the understanding of disease mechanisms and guide future strategies targeting β-cell function.

## Introduction

Loss of β-cell function is a hallmark of type 2 diabetes (T2D) (1), yet it remains unclear whether this reflects a reduction in insulin secretory granule (ISG) content or functional impairment within structurally altered cells. Ultrastructural studies based on transmission electron microscopy (TEM) have provided important insights into ISG organisation (2), but their limited sampling capacity and low scalability make large-scale quantitative analysis in human tissue challenging. Expansion microscopy (ExM) enables nanoscale imaging by physically enlarging biological specimens, allowing sub-diffraction resolution with conventional microscopes (3). In particular, Magnify, a recently developed ExM approach, is compatible with paraformaldehyde(PFA)-fixed, paraffin-embedded (PFA-PE) human tissues and preserves antigenicity for post-expansion immunostaining (4).

Here, we applied Magnify to human pancreatic tissue to quantitatively assess ISG organisation *in situ* in T2D and non-diabetic (ND) donors. We asked whether β-cell dysfunction is associated with granule depletion or instead with altered intracellular organisation. By combining nanoscale imaging with morphometric analysis, we identify increased granule clustering as a defining feature of diabetic β-cells and link this structural rearrangement to impaired insulin secretion.

## Methods

PFA-PE pancreatic tissue from ND (n=3) and T2D (n=3) donors was processed using the Magnify ExM protocol (4). 5 μm-sections were embedded in a hydrogel matrix containing methacrolein as an anchoring agent, homogenised and expanded (expansion factor 3.1 ± 0.4). Post-expansion immunostaining targeted insulin to visualise secretory granules. Confocal images were acquired under identical conditions for all samples. To quantify insulin distribution, images acquired in the 488 nm channel were first pre-processed using a bandpass filter to enhance signal-to-noise ratio. Insulin-enriched puncta were then segmented using CellPose, a deep learning-based algorithm. Morphometric parameters—including area, perimeter, and roundness—were extracted using a custom-written MATLAB script. Clusters were operationally defined as insulin-positive structures with area >0.16 μm^2^ (corresponding to the upper outlier threshold of the ND distribution, Q3 + 1.5×IQR) and roundness <0.8, consistent with the expected near-spherical morphology of insulin granule cores (5). The fraction of clustered granules with respect to the total insulin-positive spots was calculated for each donor. The number of granules per β-cell section was estimated by dividing the total insulin-positive area to the number of β-cell nuclei and to the average area of a single granule, calculated as weighted average from the bimodal granule size distribution. Donor characteristics are reported in Table 1 and included islet functional data, such as the insulin stimulation index (ISI). Full protocol details are provided in the Electronic Supplementary material (ESM).

## Results

### Expansion microscopy enables quantitative nanoscale analysis of human β-cells

Application of the Magnify protocol to PFA-PE human pancreatic tissue yielded an isotropic linear expansion of 3.1 ± 0.4, enabling nanoscale resolution imaging of insulin-labelled structures *in situ* (Figure 1a–e). The expansion factor is lower than the maximal expansion reported for Magnify and is consistent with the reduced and less uniform expansion observed in FFPE tissues(4), potentially further influenced by the architecture of the pancreas, characterized by a complex and collagen-rich extracellular matrix(6). Post-expansion imaging revealed subcellular details that were not resolvable prior to expansion, allowing robust segmentation of individual insulin-positive structures across large tissue areas (Figure 1f).

**Figure 1.**
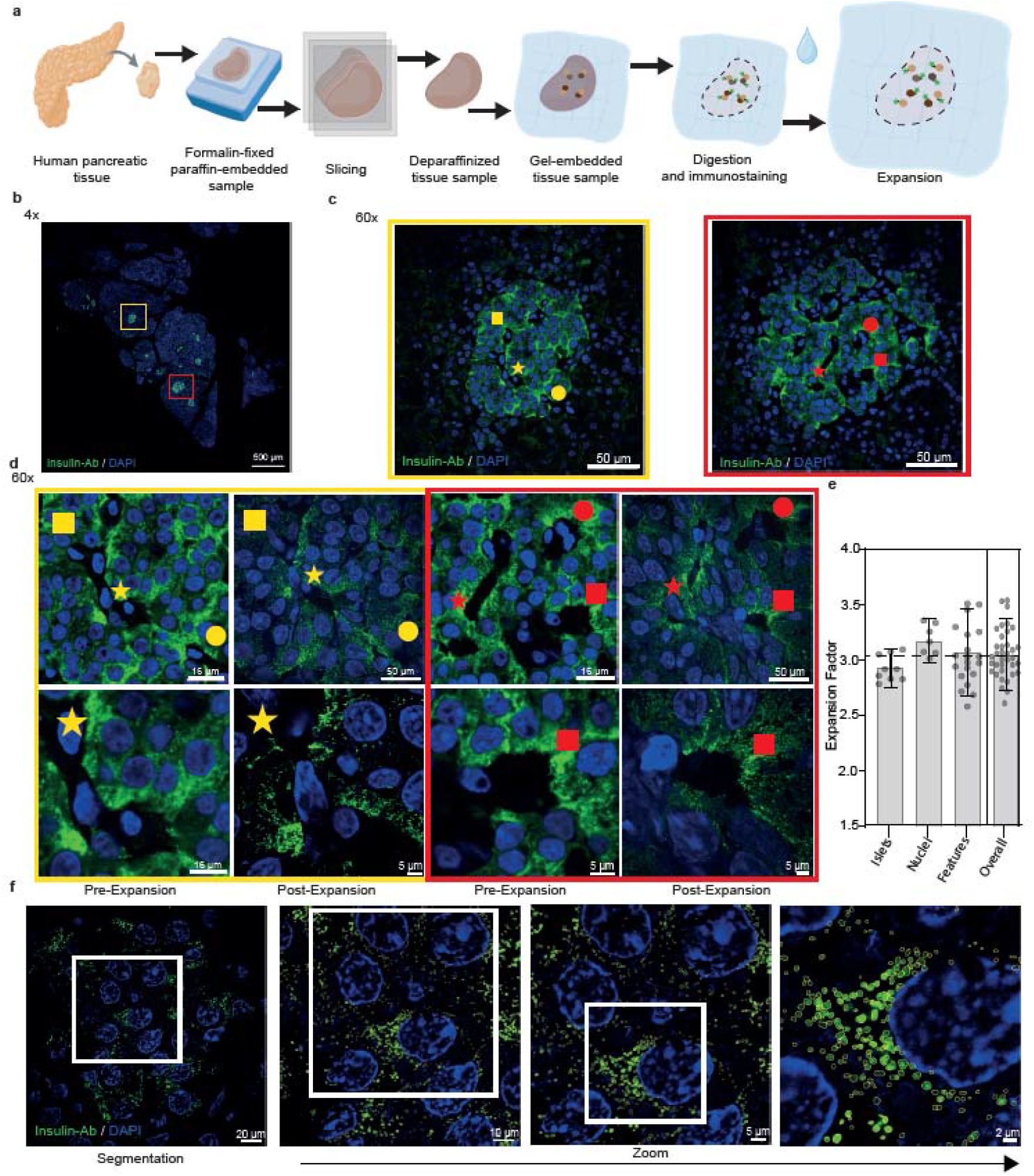
ExM enables nanoscale imaging and quantitative analysis of human pancreatic β-cells. (a) Schematic of the Magnify ExM workflow applied to PFA-PE human pancreatic tissue. (b–c) Representative pre-expansion images of human pancreatic sections stained for insulin (green) and nuclei (DAPI, blue) at low (b) and high (c) magnification, highlighting islets of Langerhans (yellow and red boxes). (d) Higher-magnification views of selected regions before and after expansion, showing improved resolution of subcellular structures. (e) Quantification of the linear expansion factor using multiple references (islets, nuclei, and structural features), yielding an average expansion of 3.1 ± 0.4. Bars represent mean ± SD. (f) Representative post-expansion images at progressively higher magnification illustrating ISG segmentation, with corresponding segmentation masks overlaid.

### Altered insulin granule organisation with preserved single-granule size and no major loss in abundance in T2D

We first assessed whether T2D is associated with changes in ISG size or morphology. Visual inspection of pancreatic tissue sections revealed differences in the spatial distribution of insulin-positive granules between ND and T2D samples (Figure 2a). Indeed, segmented insulin-positive objects in T2D were characterised by increased area and decreased roundness compared with ND samples (Figure 2b,c). To further characterize these differences, we mapped all detected structures in a 2D roundness–area space (Figure 2d). This revealed a population of larger and less circular insulin-positive structures in T2D, consistent with granule clusters as defined by combined area and roundness criteria. To analyse individual insulin granules, we applied a roundness-based filter (roundness > 0.8) to select highly circular insulin-positive objects, consistent with the expected morphology of single granules (5). The diameter distribution of this subset was similar in ND and T2D samples and showed a bimodal profile, with peaks centred at approximately 145 nm and 177 nm across donors in both groups (Figure 2e). These values are consistent with reported dense-core granule diameters measured by soft X-ray tomography(7), supporting the presence of distinct single-granule subpopulations and no major shift in individual granule size in T2D. To further assess whether β-cell alterations in T2D reflect changes in granule abundance, we estimated the number of ISGs per cell by normalizing the total insulin-positive area to the number of β-cell nuclei and to the average area of a single granule (Figure 2f). This analysis provides an approximate estimate and suggests a moderate reduction in granule number per cell in T2D samples (∼25% decrease; ND: 669.7 ± 239.1 vs T2D: 500.3 ± 162.9). Together, these results suggest changes in granule organisation in T2D, with preserved single-granule size and a moderate reduction in granule abundance.

**Figure 2.**
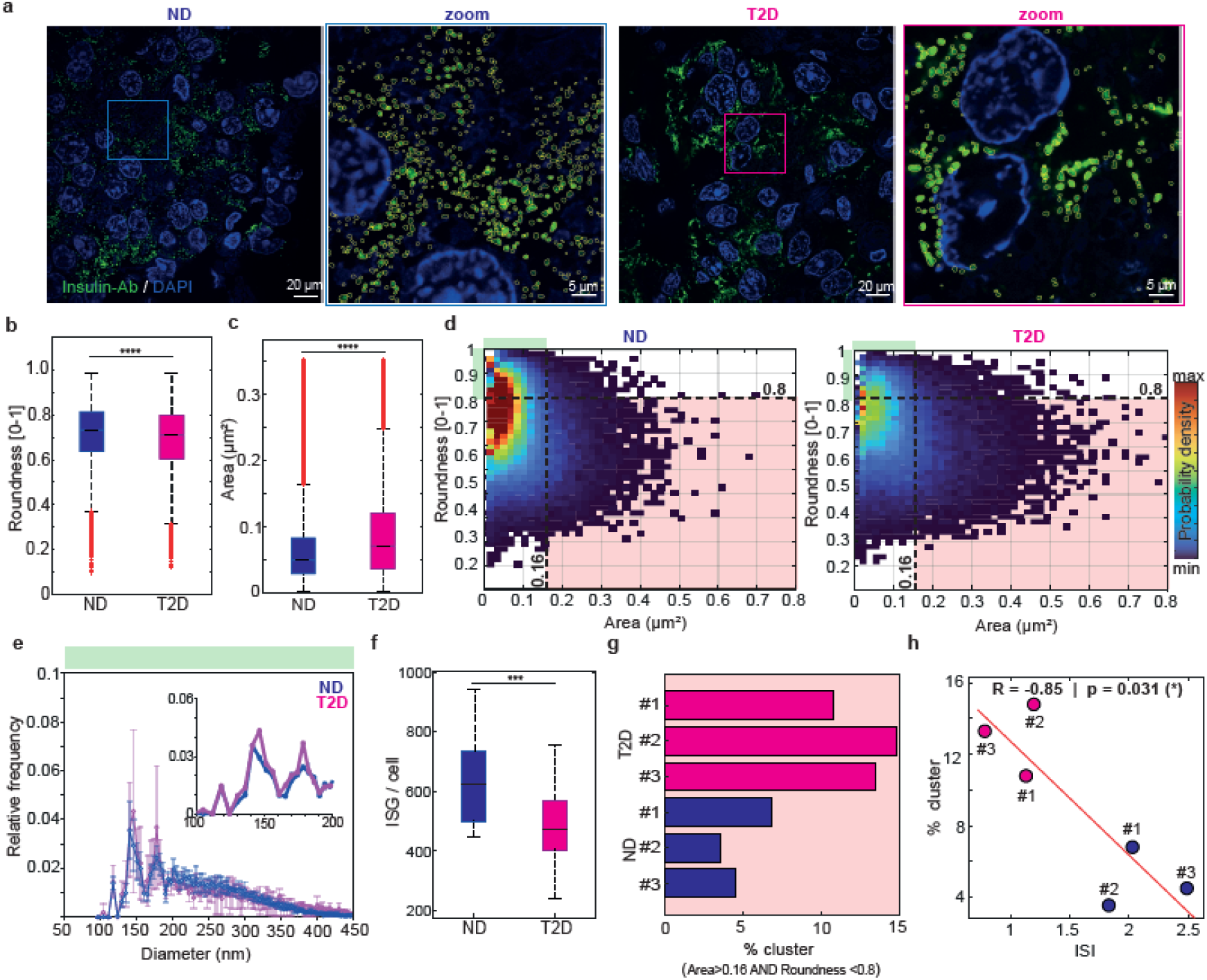
ISG organisation is altered in T2D. (a) Representative images of insulin-positive granules (green) in ND and T2D β-cells, with segmentation masks overlaid (yellow). (b–c) Quantification of granule roundness (b) and area (c) in ND and T2D samples. Data are derived from segmented insulin-positive objects (ND: n = 114,243; T2D: n = 57,285). ****p < 0.0001 (Mann–Whitney U test). (d) 2D distribution of granule roundness and area in ND and T2D. Regions corresponding to highly circular, small granules (putative isolated granules; green area) and larger, less circular structures (putative clusters; red area) are highlighted. (e) Diameter distribution of highly circular granules (roundness > 0.8), shown as mean ± SD across donors; inset highlights the bimodal region. ND and T2D samples display a similar bimodal profile. (f) Estimated number of ISGs per β-cell section (ISG/cell), calculated from the total insulin-positive area, normalized to the number of β-cell nuclei and to the average single-granule area (ND: n = 19 images from 2 donors due to failed nuclear segmentation of the third donor; T2D: n = 34 images from 3 donors). ***p < 0.001 (Mann–Whitney U test). (g) Fraction of granules classified as clusters (area >0.16 μm^2^ and roundness <0.8) in individual donors. (h) Negative correlation between granule clustering and insulin stimulation index (ISI) across all donors, assessed by Pearson’s correlation (r = –0.85, p = 0.031).

### Granule clustering is increased in T2D and correlates with impaired insulin secretion

The enrichment of larger and less circular insulin-positive structures observed in T2D (Figure 2d) points to altered granule organisation in diabetic β-cells. To quantify this effect at the donor level, we focused on the region of the roundness–area space corresponding to these structures. The fraction of clustered granules consistently increased in T2D compared with ND across all donors (Figure 2g). Importantly, granule clustering was strongly inversely correlated with ISI measured in isolated islets from the same donors (r = –0.85, p = 0.031) (Figure 2h), linking nanoscale structural reorganisation to functional impairment. Together, these observations suggest that β-cell dysfunction in T2D may not be primarily driven by a loss of ISGs, but rather by alterations in their spatial organisation.

## Discussion

By combining ExM with quantitative morphometric analysis, we establish a framework to resolve and classify insulin-positive structures directly in human tissue. Previous work from our group has applied ExM to ISGs in INS-1E cell line (8), demonstrating its potential for nanoscale analysis in simplified systems, whereas other studies have applied ExM to pancreatic islets and tissue primarily to investigate primary cilia (9,10).

Mapping granules in a roundness–area parameter space enabled discrimination between isolated granules and higher-order assemblies, providing an objective framework for quantifying clustering in situ. Our results show that β-cell dysfunction in T2D is characterized by nanoscale reorganization of ISGs rather than by their depletion. In particular, we identify increased granule clustering as a prominent feature of diabetic β-cells, which is strongly associated with impaired insulin secretion. Notably, larger and morphologically complex insulin-positive structures have previously been described as multigranular bodies associated with granule ageing and degradation pathways(11). Although our analysis does not directly establish their molecular identity, these findings raise the possibility that the increased clustering observed in T2D may, at least in part, reflect altered granule turnover or disposal.

These results support the concept that β-cell failure in T2D primarily reflects functional alterations in surviving cells rather than a reduction in ISG content. While ultrastructural validation will be important in future studies, the large-scale quantitative analysis enabled by ExM provides a complementary approach to assess granule organization in human tissue.

Overall, our findings highlight ExM as a powerful approach for nanoscale analysis of human pancreatic tissue. Future studies integrating additional molecular markers will be important to define the mechanisms underlying granule clustering and to enable multiplexed nanoscale analysis of endocrine and exocrine compartments within pancreatic tissue.

## Acknowledgements

We thank Dr Aleksandra Klimas for technical support with Magnify.

## Contribution statement

Conceptualisation: FC, PM, YZ, VDL; Methodology: AL, YZ, VDL, GF; Investigation: LAP, VDL, HV, MT, CDL, MS; Data curation: LAP, GF, VDL; Formal analysis: LAP, VDL; GF, MT; Visualization: GF; Resources: LM, PM, YZ, FC, FB; Supervision: FC; Funding acquisition: FC; Writing – original draft: VDL, FC; Writing – review & editing: VDL, FC, GF, PM, YZ, FB. All authors approved the final manuscript.

## References

1. Yau B, Ghislain J, Kebede MA, Hughes J, Poitout V. The role of the beta cell in type 2 diabetes: new findings from the last 5 years. Diabetologia. 2025;68(10):2092–103.

2. McLaughlin MR, Weaver SA, Syed F, Evans-Molina C. Advanced Imaging Techniques for the Characterization of Subcellular Organelle Structure in Pancreatic Islet β Cells. Compr Physiol. 2023;14(1):5243–67.

3. Wassie AT, Zhao Y, Boyden ES. Expansion microscopy: principles and uses in biological research. Nat Methods. 2019;16(1):33–41.

4. Klimas A, Gallagher BR, Wijesekara P, Fekir S, DiBernardo EF, Cheng Z, et al. Magnify is a universal molecular anchoring strategy for expansion microscopy. Nat Biotechnol. 2023;41(6):858–69.

5. Zhang X, Peng X, Han C, Zhu W, Wei L, Zhang Y, et al. A unified deep-learning network to accurately segment insulin granules of different animal models imaged under different electron microscopy methodologies. Protein Cell. 2019;10(4):306–11.

6. Santos da Silva T, Silva-Júnior LN da, Horvath-Pereira B de O, Valbão MCM, Garcia MHH, Lopes JB, et al. The Role of the Pancreatic Extracellular Matrix as a Tissue Engineering Support for the Bioartificial Pancreas. Biomimetics. 2024 Oct 2;9(10):598.

7. Deshmukh A, Chang K, Cuala J, Campos MJH, Mahmood S, Verma R, et al. Secretory stimuli distinctly regulate insulin secretory granule maturation through structural remodeling. Structure. 2025;33(11):1831-1843.e4.

8. Pugliese LA, Lorenzi V De, Bernardi M, Ghignoli S, Tesi M, Marchetti P, et al. Unveiling nanoscale optical signatures of cytokine-induced β-cell dysfunction. Sci Rep. 2023;13(1):13342.

9. Müller A, Klena N, Pang S, Garcia LEG, Topcheva O, Duran SA, et al. Structure, interaction and nervous connectivity of beta cell primary cilia. Nat Commun. 2024;15(1):9168.

10. Dong X, Jo JH, Hughes J. Axonemal Dynein Visualized in Primary Cilia via Expansion Microscopy. Cytoskeleton. 2025;1–13.

11. Hoboth P, Müller A, Ivanova A, Mziaut H, Dehghany J, Sönmez A, et al. Aged insulin granules display reduced microtubule-dependent mobility and are disposed within actin-positive multigranular bodies. Proc Natl Acad Sci U S A. 2015;112(7):E667–76.

