## Supplementary information for "Expansion microscopy reveals insulin granule clustering in human β-cells in type 2 diabetes"

**Electronic Supplementary Material – Extended Methods**

**Human pancreatic tissue samples**

Pancreases were obtained from heart-beating non-diabetic (ND, n=3) and type 2 diabetic (T2D, n=3) organ donors received before November 2021 and processed with the approval of the Ethics Committee of the University of Pisa (21 November 2013, #2615). Table 1 report the main clinical characteristics of the organ donors and isolated islet insulin secretion features. Pancreatic samples, collected from the neck of the pancreas, were fixed in 4% PFA overnight at 4°C and embedded in paraffin. Haematoxylin-eosin (HE) staining was performed on 2 μm thick-section to evaluate the quality of the tissue (adequate and uniform fixation, preservation of tissue structures, absence of artefacts).

| **ND/T2D** | **SEX** | **AGE (years)** | **BMI (Kg/m2)** | **Cause of death** | **Insulin Release (uU/islet/min) at 3.3 mM glucose** | **Insulin Release (uU/islet/min) at 16.7 mM glucose** | **Insulin Stimulation Index (ISI)** |
| --- | --- | --- | --- | --- | --- | --- | --- |
| ND | F | 82 | 27.6 | Cardiovascular disease | 0.300 | 0.581 | 2.03 |
| ND | M | 75 | 23.2 | Cardiovascular disease | 0.05 | 0.12 | 2.49 |
| ND | F | 77 | 31.3 | Cardiovascular disease | 0.197 | 0.371 | 1.83 |
| T2D | M | 81 | 26.4 | Cardiovascular disease | 0.042 | 0.39 | 0.80 |
| T2D | F | 71 | 31.1 | Cardiovascular disease | 0.037 | 0.052 | 1.16 |
| T2D | F | 68 | 28.3 | Cardiovascular disease | 0.079 | 0.1 | 1.22 |

**Table 1: Main clinical characteristics of the organ donors and isolated islet function.**

**Islet purification and insulin secretion assay**

Islets were isolated from the same donors used for pancreatic tissue analysis. Islet isolation was performed by collagenase digestion followed by density gradient purification. Isolated islets were cultured at 37°C in a humidified atmosphere with 5% CO₂ in M199 medium supplemented with 10% (v/v) adult bovine serum, 100 U/mL penicillin, 100 μg/mL streptomycin, 750 ng/mL amphotericin B and 50 μg/mL gentamicin, at 5.5 mM glucose, as previously reported(1). For insulin secretion assays, batches of 15 handpicked islets were pre-incubated for 45 min in Krebs’ solution (KRBS; 115 mM NaCl, 5 mM KCl, 24 mM NaHCO_3_, 1 mM MgCl_2_, 2.5 mM CaCl_2_, 0.5% BSA, pH 7.4) containing 3.3 mM glucose, followed by static incubation for 45 min at 3.3 or 16.7 mM glucose. Supernatants were collected and stored at −20°C. Insulin levels were measured by ELISA (Mercodia AB, Uppsala, Sweden) according to the manufacturer’s instructions. The insulin stimulation index (ISI) was calculated as the ratio of insulin secretion at 16.7 mM glucose to that at 3.3 mM glucose.

**Magnify expansion microscopy**

The Magnify protocol was performed as described by Klimas et al.(2), with minor modifications to accommodate PFA-PE tissues. PFA-PE blocks were sectioned at 5 μm using fully automated rotary microtome (RM2255, Leica Microsystems, Wetzlar, Germany). Sections were baked at 60°C for 1 h, deparaffinized in xylene (10 min), rehydrated through graded ethanol (100%, 95%, 75%, 50%; 10 min each at room temperature) and rinsed in distilled water. A monomer solution containing sodium acrylate (34% w/v), acrylamide (10% w/v), N,N-dimethylacrylamide (4% v/v), NaCl (1% w/v), and bisacrylamide (0.01% w/v) in PBS was prepared. Polymerisation was initiated by addition of ammonium persulfate (APS, 0.25% w/v), TEMED (0.1% v/v), 4-hydroxy-TEMPO (4-HT, ~0.001%) and methacrolein (0.05% v/v) as anchoring agent. A drop of the polymerisation solution was applied to fully cover the tissue section, followed by incubation at 4°C for 30 min to allow diffusion. Gelation was then carried out at 37°C overnight in a humidified chamber. Following gelation, excess gel surrounding the tissue was removed and the gel-embedded sections were transferred to microcentrifuge tubes. Samples were incubated in a homogenisation buffer containing SDS (10% w/v), 8 M urea, 25 mM EDTA, 0.1 M glycine, 0.5 M Tris and PBS (2×), adjusted to pH 8.0. Homogenisation was performed by heat treatment in a pressure cooker at 12 psi for 20 min. Subsequently, gels were washed in a surfactant solution (1% C12E10 in PBS supplemented with sodium azide). Samples were washed three times for 10 min at room temperature, followed by a 1 h wash at 60°C, and subsequently washed three additional times in the same buffer to ensure complete removal of residual reagents. Gels were incubated with anti-insulin primary antibody (Immunological Sciences #AB-84377, diluted 1:500) in 0.2% surfactant solution at 37°C overnight with gentle agitation. Following three washes (10 min each) in 0.2% surfactant solution, samples were incubated with anti-rabbit IgG Alexa Fluor 488 (Immunological Sciences #IS20015-1, diluted 1:500) and DAPI (ThermoFisher # 62248, diluted 1:1000) in the same buffer for 3 h at 37°C. Gels were then washed twice for 10 min in 0.2% surfactant solution and once in PBS.

**Expansion factor estimation**

For estimation of the expansion factor, a dedicated sample was immunostained prior to gelation. Following deparaffinisation and rehydration, antigen retrieval was performed in 20 mM sodium citrate buffer (pH 8.0) at 60°C for 30 min, followed by cooling to room temperature and washing in PBS. Sections were then blocked in 5% BSA in PBS containing 0.1% Tween for 1–1.5 h at room temperature and incubated with anti-insulin primary antibody followed by secondary antibody. Samples were then processed through gelation, homogenisation and expansion according to the Magnify protocol described above. The expansion factor was estimated by comparing corresponding features in pre- and post-expansion images across multiple structures.

**Imaging**

Fluorescence imaging was performed using a Nikon Eclipse Ti2 microscope equipped with a CSU-W1 spinning disk confocal module and a Hamamatsu Fusion sCMOS camera (C14440-20UP), controlled by NIS-Elements AR software (v. 5.21.03). Images were acquired using the following Nikon objectives: CFI Plan Apo Lambda ×10 (0.45 NA, air), CFI Apo LWD Lambda S ×20 (0.95 NA, water), CFI Apo LWD Lambda S ×40 (1.15 NA, water) and Plan Apo VC 60×A WI DIC N2 (1.2 NA, water). Excitation was provided by 405 nm and 488 nm laser lines for DAPI and insulin, respectively, and emission was collected using DAPI and FITC emission filters. Images were acquired at 16-bit depth with 1×1 binning. For 60× imaging, the pixel size was at ~100 nm. Acquisition settings were kept constant across all samples.

**Image processing and quantitative analysis**

Image analysis was performed using Fiji (ImageJ) and MATLAB. Images acquired in the 488 nm channel were first pre-processed with a bandpass filter to improve signal-to-noise ratio. Insulin-positive structures were then segmented using CellPose. The pretrained model “cyto” was used to segment both ND and T2D images, using 6 pixel diameter as starting value. Cell probability threshold parameter was set to the default value and flow threshold parameter was set to 0.8. Morphometric parameters, including area, perimeter and roundness, were extracted from the segmented masks using a custom-written MATLAB script.

*ISG size analysis*

To analyse putative single granules, we selected highly circular insulin-positive objects (roundness >0.8), consistent with the expected near-spherical morphology of individual granules(3). Granule diameter distributions were calculated from the segmented objects after correction for the expansion factor.

*ISG clustering*

Clusters were operationally defined as insulin-positive structures with area >0.16 μm² (threshold derived from the upper outlier threshold of the ND distribution: Q3 + 1.5×IQR) and roundness <0.8, corresponding to the population of larger and less circular objects identified in the roundness–area space. The fraction of clustered granules was calculated for each donor as the percentage of insulin-positive structures falling within this region relative to the total number of segmented objects.

*ISG number estimation*

The number of granules per β-cell section was estimated by normalising the total insulin-positive area to the number of β-cell nuclei and to the average area of a single granule. The latter was calculated from the bimodal granule size distribution by computing the mean diameter of each population and deriving a weighted average based on their relative frequencies. β-cell nuclei were identified based on insulin positivity of the surrounding cytoplasm; this approach provides an approximate estimate of granule abundance per β-cell section.

**Methods References**

1.	Suleiman M, Sawatani T, Tesi M, Yi X, Papadopoulou T, Rufer C, et al. Functional recovery of islet β cells in human type 2 diabetes : Transcriptome signatures unveil therapeutic approaches. Sci Adv. 2025;11(41):eads2905.

2.	Klimas A, Gallagher BR, Wijesekara P, Fekir S, DiBernardo EF, Cheng Z, et al. Magnify is a universal molecular anchoring strategy for expansion microscopy. Nat Biotechnol. 2023;41(6):858–69.

3.	Zhang X, Peng X, Han C, Zhu W, Wei L, Zhang Y, et al. A unified deep-learning network to accurately segment insulin granules of different animal models imaged under different electron microscopy methodologies. Protein Cell. 2019;10(4):306–11.
